# Limiting Brownian Motion to Enhance Immunogold Phenotyping and Superimpose Optical and Non-Optical Single-EP Analyses

**DOI:** 10.1101/2024.02.22.581663

**Authors:** Kim Truc Nguyen, Xilal Y. Rima, Colin L. Hisey, Jacob Doon-Ralls, Chiranth K. Nagaraj, Eduardo Reátegui

**Author notes:** Authors contributed equally (Kim Truc Nguyen, Xilal Y. Rima).

## Abstract

Optical and non-optical techniques propelled the field of single extracellular particle (EP) research through phenotypic and morphological analyses, revealing the similarities, differences, and co-isolation of EP subpopulations. Overcoming the challenges of optical and non-optical techniques motivates the use of orthogonal techniques while analyzing extracellular particles (EPs), which require varying concentrations and preparations. Herein, we introduce the nano-positioning matrix (NPMx) technique capable of superimposing optical and non-optical modalities for a single-EP orthogonal analysis. The NPMx technique is realized by ultraviolet-mediated micropatterning to reduce the stochasticity of Brownian motion. While providing a systematic orthogonal measurement of a single EP via total internal reflection fluorescence microscopy and transmission electron microscopy, the NPMx technique is compatible with low-yield samples and can be utilized for non-biased electrostatic capture and enhanced positive immunogold sorting. The success of the NPMx technique thus provides a novel platform by marrying already trusted optical and non-optical techniques at a single-EP resolution.

## Introduction

Extracellular particles (EPs) are highly heterogeneous with various particle subpopulations, including lipoproteins (LPs), extracellular vesicles (EVs), viruses, and non-vesicular EPs.^1^ With the increasing interest in EVs, which are cell-derived lipid nanoparticles that mediate aspects of interorgan communication^2-4^ and are biomolecular fountains for liquid biopsies,^5-7^ their isolation and subsequent characterization have been paramount. However, EPs overlap in size and density; thus, their isolation is confounded by co-isolates of differing EP subpopulations.^8-10^ Therefore, the field has encouraged using orthogonal methods to ensure EP purity.^11^ Traditional molecular techniques for the characterization of EPs require the bulk lysis of particles and dilute co-isolate profiles, promoting single-EP methods to investigate the degree of co-isolation or heterogeneity within EP subpopulations. Recently, single-EV detection utilizing optical and non-optical microscopic techniques has identified the co-isolation of EVs and LPs at the single-EP level,^12-15^ emphasizing the motivation to characterize EPs at a single-EP resolution to comprehend co-isolation dynamics. However, both techniques have limitations. Super-resolution microscopy (SRM) has demonstrated potential in clarifying the heterogeneity of EPs through fluorescence co-localization and subpopulation-based positive immunoselection.^15^ However, EPs that downregulate the epitopes targeted by the fluorescent probes are excluded from the analyses thus producing false-negative biases. Furthermore, morphologies of the particles are obscured due to the diffraction limit.^16^ On the other hand, while electron microscopy (EM) has provided fundamental morphological data on EPs and visualizes all EPs without biases to antigens, phenotyping via immunogold labeling is hardly quantifiable due to the low efficiencies and limitations in multiplexing.^17^ Therefore, a technique that provides both the phenotyping ability of SRM techniques and the morphological quantification of EM techniques would further enhance the field and limit biases in quantification.

Herein, we introduce the nano-positioning matrix (NPMx) technique capable of performing orthogonal measurements at the single-EP resolution via superimposing micropatterns acquired through non-optical and optical microscopy. Micropatterning on transmission EM (TEM) grids through ultraviolet (UV)-induced degradation of a non-biofouling polymer brush enables the precise functionalization of the mesh surface in traceable coordinates. The tunable design provides electrostatic entrapment of EPs for a non-biased capture of the entire subpopulation and facile immunogold labeling by limiting Brownian motion via tethering antibody-functionalized gold nanoparticles (AuNPs) onto the mesh surface. The distinctive micropatterns enabled the one-to-one mapping of SRM images to EM images to correlate fluorescence signals to EVs captured on the mesh surface. The engineered surface thus allows for a systematic technique that enhances EM efficiencies, improves immunogold labeling, and provides a framework for an orthogonal measurement of the same EP.

## Materials and Methods

### Fluorescent EV collection

Gli36 cells were transfected with PalmtdTomato^18,19^ and were maintained in Dulbecco’s Modified Eagle Medium (DMEM) supplemented with 10 % (v/v) fetal bovine serum (FBS) and 1 % (v/v) penicillin-streptomycin (PS). Once 70 % confluence was achieved, the cells were washed with phosphate-buffered saline (PBS) and given serum-free DMEM for 48 hours after which the supernatant was collected and centrifuged at 2,000 ×*g* for 10 min to remove cell debris. The supernatant was then purified via size-exclusion chromatography (SEC) with a 70 nm qEV original column (Izon Science) by pooling EV-rich fractions 1-4, according to the manufacturer’s instructions.

### Bioreactor EV collection

As previously described, U373 cells were cultured in a CELLine AD 1000 adherent bioreactor flask (Wheaton).^20,21^ Briefly, 25 × 10^6^ cells were inoculated in 15 mL of complete DMEM with 10 % FBS and 1 % PS in the lower cell chamber with ∼750 mL of the same media added to the upper media chamber. The bioreactor was then gradually adapted from 10 % FBS supplementation to 10 % chemically-defined CDM-HD (Fibercell) serum replacement over four weeks to avoid exogenous FBS EP contamination. EVs were collected in ∼15 mL of conditioned media three times per week from the cell chamber, and the media chamber was refreshed once weekly. The 15 mL of conditioned media was centrifuged at 2,000 ×*g* for 10 min to remove suspended cells and large debris, and the resulting supernatant was concentrated using a 100 kDa MWCO Vivaspin 20 filter (Sartorius) at 3,000 ×*g* to a volume of ∼4 ml. This concentrated media was centrifuged at 10,000 ×*g* for 30 min to remove large EVs, and the remaining supernatant was further concentrated using a 100 kDa Vivaspin 500 filter (Sartorius). The crude small EV suspension was then purified with SEC as previously described.

### Tunable resistive pulse sensing (TRPS)

Tunable resistive pulse sensing (qNano Gold, Izon Science) was utilized to measure the concentration and size of EVs using a stretchable NP200 membrane. A pressure of 0.5 kPa and voltage of 0.38 V were maintained throughout the measurements. The measurements were calibrated with known standard polystyrene nanoparticles of 200 nm (CPC200, Izon Science).

### TEM grid micropatterning

The working volume utilized for an individual grid was 20 μL, which was kept constant for the different solutions added onto the grids. Various TEM grids were used in this investigation, including Formvar/Carbon 75 mesh, Copper (Ted Pella, cat# 01802-F), Carbon Type-B, 200 mesh TH, Nickel (Ted Pella, cat# 01808N), Carbon Type-B, 300 mesh, Nickel (Ted Pella, cat# 01813N). The TEM grids were primed with 70 % (v/v) ethanol and quickly transferred to PBS via three washes. The grids were then transferred to a 100 µg/mL solution of poly-L-lysine-*graft*-poly(ethylene glycol) (PLL-*g*-PEG) and incubated for 1 hour. The excess PLL-*g*-PEG was washed away by transferring the functionalized grid to PBS three times. After that, PLPP, a commercial photoactivator (Alvéole, France), was transferred onto the functionalized grid. The grid was then subjected to UV-induced degradation of the PLL-*g*-PEG by the PRIMO micropatterning system (Alvéole, France). After micropatterning, the grids were transferred to PBS and washed three times to remove the remaining PLPP.

### Electrostatic capture of EPs

A 0.1 % (w/v) poly-L-lysine (PLL) solution was incubated on the TEM grid post-micropatterning for 5 min to allow its adsorption onto the micropatterned regions. The grids were then transferred to PBS and washed three times. The EV solution was then incubated onto the functionalized TEM grid for 1 min whereby the unbound EVs were subsequently washed off with PBS by pipetting 10 times in an up-and-down motion. The rinsing step was repeated 10 times.

### Gold nanoparticle immobilization

A solution of 50-nm streptavidin-functionalized AuNPs (Strep-AuNP) at various concentrations, including 0.25 % and 0.5 % (w/v) (GNA50, Nanocs Inc.), was incubated onto the micropatterned TEM grid for 15 min. The grids were then washed with PBS by pipetting PBS 10 times in an up-and-down motion. This rinsing step was repeated 10 times.

### Immunogold capture of EPs

TEM grids functionalized with Strep-AuNPs were blocked with a 3 % (w/v) of bovine serum albumin (BSA) solution (Sigma-Aldrich) diluted in PBS for 15 min to reduce non-specific capture. Capture antibodies (provided in **Supplementary Table 1**) were biotinylated using the EZ-Link micro Sulfo-NHS-biotinylation kit from (Thermo Fisher Scientific). The biotinylated antibodies were then diluted to 10 μg/mL in a 1 % BSA solution and added to the grids for 15 min. Excess antibodies were rinsed away by pipetting PBS 10 times in an up-and-down motion. The rinsing step was repeated 10 times. The surface was further blocked with a 3 % (w/v) solution of BSA for 15 min. After removing the BSA solution, murine leukemia virus tagged with V5 (MLV-V5, ViroFlow Technologies) or EVs were subsequently incubated onto the grids for 40 min at room temperature in a humid environment. The unbound virions and EVs were washed away by pipetting PBS 10 times in an up-and-down motion. The rinsing step was repeated 10 times.

### SRM image acquisition

Total internal reflection fluorescence microscopy (TIRFM; Nikon, Melville, NY) was utilized to acquire SRM images. The functionalized TEM grids were placed onto a coverslip and imaged via TIRFM with a 100 × objective and immersion oil to reduce surface refraction. The functionalized TEM grids were maintained in PBS throughout the TIRFM imaging process.

### Negative staining

Two 20 µL droplets of water for injection (WFI) and two 20 µL droplets of negative stain (UranyLess EM stain, Electron Microscopy Sciences) were placed onto parafilm. The functionalized TEM grids were then immersed successively into the two WFI droplets and carefully blotted to remove excess liquid. The grids were then stained via submersion into a droplet of the negative stain, blotted, submersed into the second droplet for ∼22 s, and then gently wicked away using filter paper. The stained TEM grids were stored in a grid box overnight to ensure thorough drying.

### EM image acquisition

TEM brightfield imaging was used at all magnifications. A low magnification range mode at 300× was utilized to locate the center and image the micropatterns. A high magnification range over the selected area at an aperture mode of 15500× and spot size 3 was used for enhanced image quality to view the lower-contrast EPs. Serial acquisition was performed with the SerialEM software (Nexperion, Austria) for a 10×10 array of images. A montage was generated with an Orius SC200 camera via binning 1, pixel size 1.84 nm, number of pieces in X × Y = 10 × 10, piece size in X × Y = 2048 × 2048 pixels, overlap in X × Y = 206 × 206 pixels, and a total area of 34.3 × 34.3 µm.

## Results

Micropatterning in TEM grids has been performed to control the spatial distribution of cells onto the mesh to maximize cellular events.^22^ Therefore, we considered a similar approach to immobilize EPs onto TEM grids to enhance the probability of EPs on the TEM-mesh surface. Briefly, the TEM grids were functionalized with PLL-*g*-PEG, which forms a brush on the surface and inhibits the adsorption of protein species. After that, the polymer brush was photoetched in specific coordinates utilizing a digital-micromirror device (DMD)-based ultraviolet (UV) projection system to induce the adsorption of protein species (**Fig. 1a**).^23^ Asymmetric micropatterns, in our case single-digit numerals, were utilized to determine the orientation of the micropatterns and facilitate superimposition of microscopic techniques (**Fig. 1b**). The DMD-based micropatterning further enabled the tunability of micropatterns insofar as single-digit or multi-digit arrays in the mesh surface for the adsorption of proteins or protein-conjugated nanoparticles (**Fig. 1c, Supplementary Fig. 1**). Immunogold labeling is riddled with bottlenecks, including the stochasticity of particle collision in a colloid mixture due to Brownian motion and the low efficiencies of labeled EPs on the mesh surface post-sample preparation.^17,24^ To address these bottlenecks, we proposed immobilizing Strep-AuNPs onto the micropatterns for their subsequent functionalization with antibodies targeting epitopes on EPs (**Fig. 1d**). The strategy thus eliminates the Brownian motion of a particle subset in the colloid mixture, in this case the Strep-AuNPs, improving capture and immobilizing the EPs onto the mesh surface and increasing the probability of a labeled EP in the field of view. Given the distinct features of the asymmetric micropattern that SRM and EM visualize, the images can be easily superimposed to correlate fluorescent signals to labeled EPs on the mesh surface (**Fig. 1e**). We term the technique of superimposing SRM and EM images as the NPMx technique.

**Figure 1:**
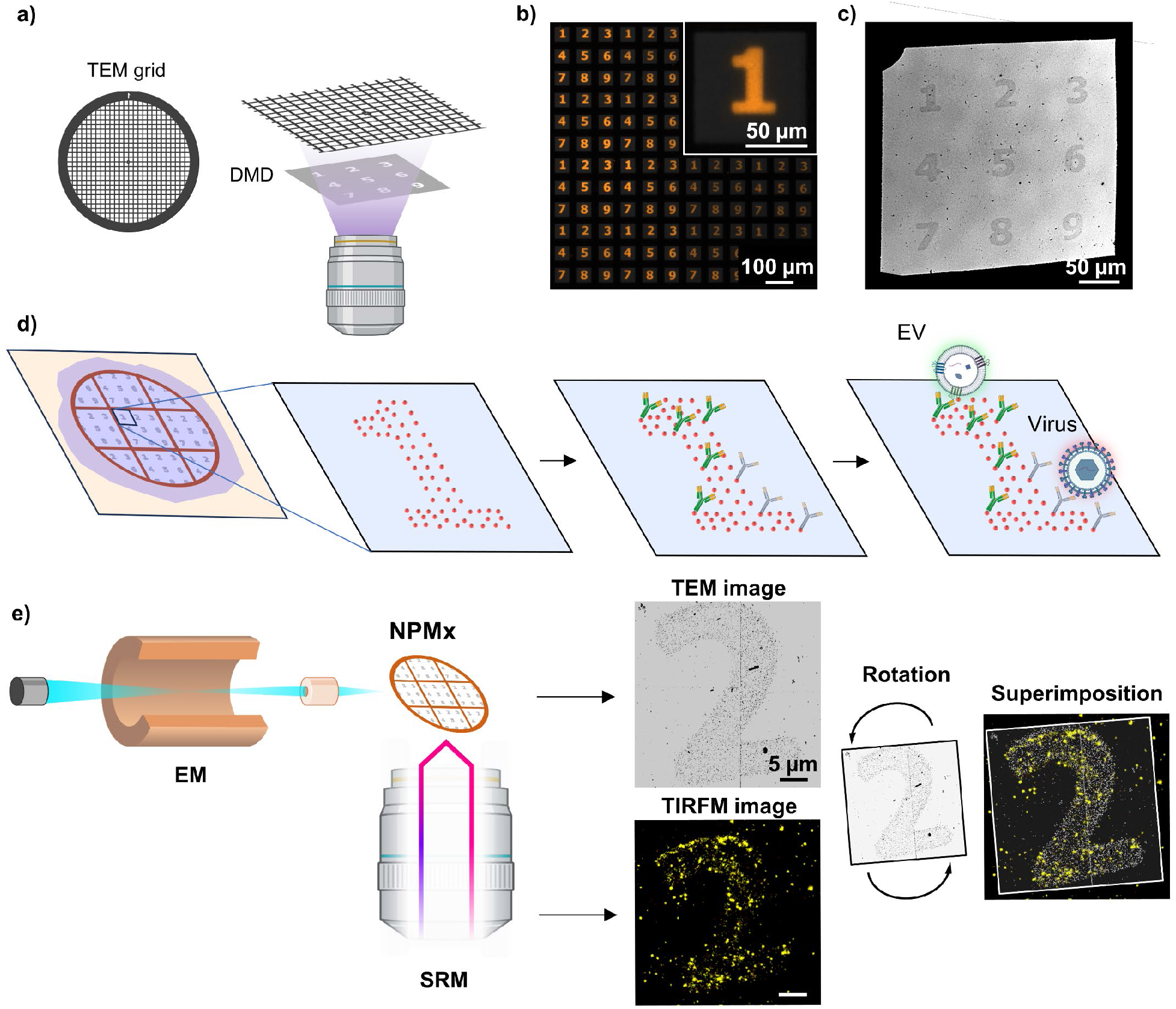
NPMx technique to superimpose optical and non-optical images at the single-EP resolution. **(a)** Transmission electron microscopy (TEM) grids are functionalized with a non-biofouling polymer film, which is degraded in specific surface coordinates mediated by a digital-micromirror (DMD)-based ultraviolet (UV) projection system. **(b)** Epifluorescence images of the photoetched micropatterns demonstrate the adsorption of protein species as asymmetrical numerals. **(c)** TEM images provide evidence for the binding of 50-nm streptavidin-functionalized gold nanoparticles (Strep-AuNP) onto the micropatterns. **(d)** The micropatterns thus enable the subsequent functionalization with biotinylated antibodies for positive immunogold sorting. **(e)** The distinctive patterns visualized by fluorescent super-resolution microscopy (SRM) and electron microscopy (EM) enable the one-to-one mapping of optical and non-optical images via rotation and superimposition.

To test the feasibility of the micropatterning method to tether EPs onto the mesh surface, EPs were directed towards the micropatterns via electrostatic interactions to induce the capture of all EPs without capture biases. Therefore, PLL, a polycation, was adsorbed onto the photoetched regions to attract negatively charged EVs. SEC-purified EVs from a glioma cell line bioreactor culture^20,21^ at a concentration of ∼2 × 10^10^ EVs/mL were added to the PLL surface demonstrating the capture of a heterogeneous population of EVs strictly onto the micropatterned region (**Fig. 2a**). To scan the entire mesh surface and visualize the numeral, a 10 × 10 array of TEM images were montaged via SerialEM, a software enabling automation of large-scale EM acquisition.^25^ Within the subpopulation of EVs, various morphologies were observed including, classical “cup-shaped” EVs, electron-dense EVs, multicomponent EVs, membrane-compromised EVs, and minuscule EP structures (**Supplementary Fig. 2**). The size distribution revealed a bimodal distribution, including an ∼11-nm sized normally distributed EP and a small EV subpopulation (**Supplementary Fig. 3**). Interestingly, TRPS measurements obscured the bimodal distribution due to the size range falling below the lower limit of detection (**Supplementary Fig. 4**).

**Figure 2:**
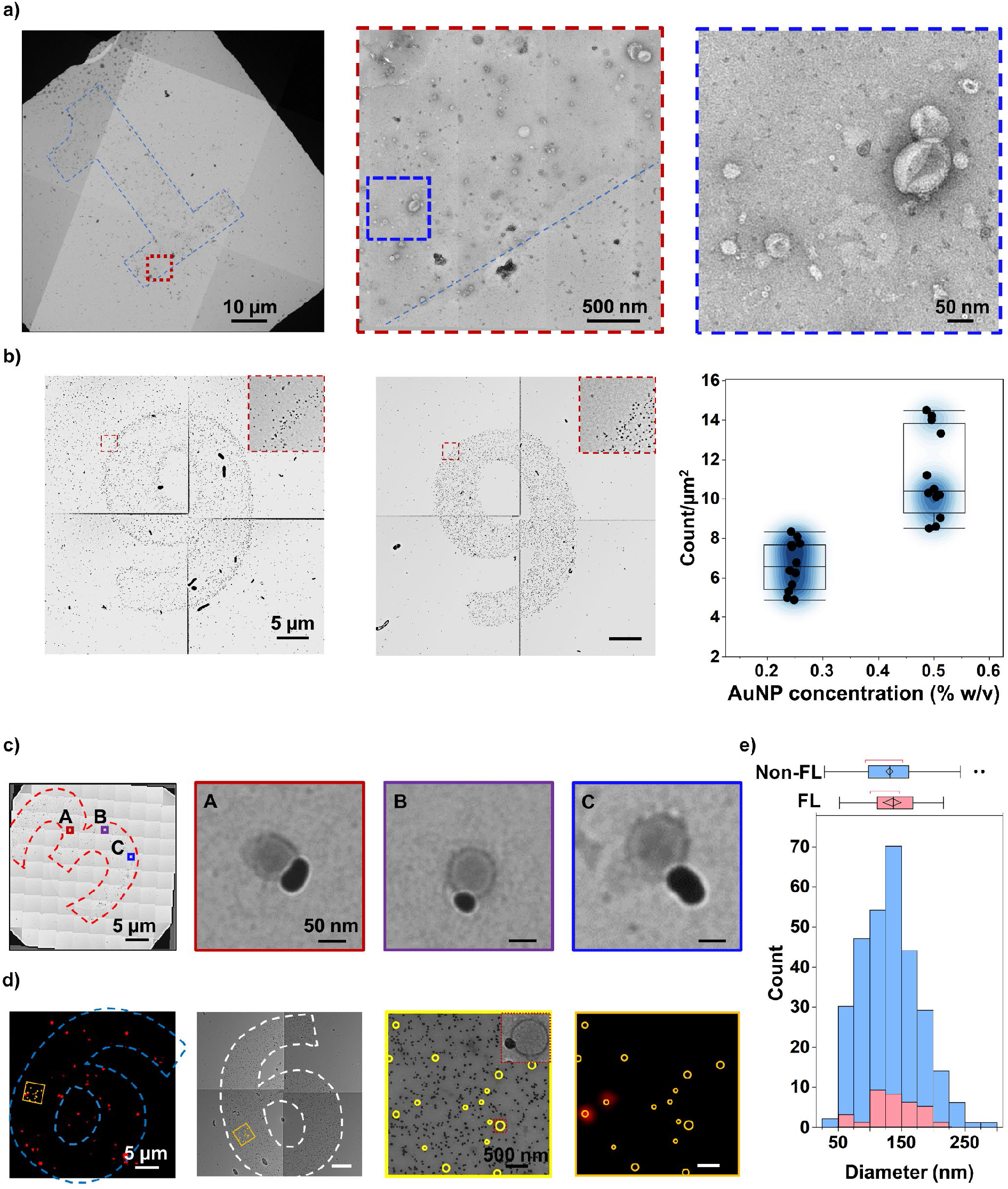
Adaptability of the NPMx technique to reduce colloidal stochasticity. **(a)** The electrostatic capture of extracellular particles (EPs) with a polycation-functionalized micropattern enables the non-biased immobilization of EPs. The dotted squares are magnified in the proceeding insets. **(b)** The adsorption of Strep-AuNPs onto the micropatterned surface is concentration-dependent, allowing for tunability in design (N = 3 numerals, n = 4 fields of view). The dotted squares are magnified in the insets. The calculations are provided in **Supplementary Table 2-5. (c)** Antibody-functionalized Strep-AuNPs facilitate immunogold labeling of murine leukemia viruses (MLVs) onto the TEM grid mesh surface. The dotted outline represents the boundary of the micropattern. The three squares are magnified in the proceeding insets. **(d)** Total internal reflection fluorescence microscopy (TIRFM) and TEM are mapped one-to-one allowing the correlation of non-optical and optical images of single extracellular vesicles (EVs). The dotted outlines represent the boundary of the micropattern. The squares are magnified in the proceeding insets. On the magnified TEM image, the square is magnified as an inset and the circles define the perimeters of the EVs. **(e)** The distribution of fluorescent (FL) and non-FL particles are represented as a histogram-box-plot conjugate, demonstrating the similarity between non-FL and FL EVs as measured in the TEM images (N = 4 numerals, n = 3 fields of view).

To improve immunogold labeling and demonstrate the NPMx technique, 0.25 % (w/v) and 0.50 % (w/v) of Strep-AuNPs were added to the photoetched regions. Similar to the two-fold increase in concentration, the Strep-AuNPs adsorbed onto the surface normalized by the surface area increased from 6.62 ± 1.24 to 11.20 ± 2.22 particles/µm^2^ (Student’s one-tailed *t-*test, *p* < 0.0001), indicating concentration-dependent adsorption of the nanoparticles to the mesh surface (**Fig. 2b, Supplementary Fig. 2-4**). To prove the concept of immunogold labeling, the Strep-AuNPs were functionalized with antibodies targeting a V5 tag, which is a short peptide sequence frequented in protein purification.^26^ MLV virions engineered to express the V5 tag were incubated on the surface at a concentration of ∼7 × 10^9^ virions/mL. As suspected, the MLV virions were observed in proximity to the functionalized AuNPs with little non-specificity outside the micropattern region (**Fig. 2c**). In the absence of an antibody, the EPs were unable to bind to the micropatterned surface, thus ensuring that binding events were a result of antibody-epitope interactions (**Supplementary Fig. 5**).

Having a workflow that enabled the immunogold immobilization of EPs in traceable surface coordinates, the NPMx technique was utilized to image the same grid via SRM and EM in an effort to map the micropatterns from one modality to the other. Therefore, EVs engineered to fluoresce via the cellular transfection of PalmtdTomato^18,19^ were added to Strep-AuNPs functionalized with antibodies targeting CD63. TIRFM images demonstrated the enrichment of fluorescent signals originating from the micropatterns and forming distinguishable numerals, indicating the capture of EVs within the micropattern with minimal non-specificity outside the micropatterned region. Similarly, EM images yielded distinct numerals via the uniform distribution of AuNPs across the micropatterned surface with EVs binding to the AuNPs. Therefore, the numerals in SRM and EM images were rotated to match with an in-house algorithm, allowing for the one-to-one mapping of captured EVs to fluorescent signals post-offset correction (**Fig. 2d, Supplementary Fig. 6a**). To our surprise, a majority of the EVs were not fluorescent (Welch’s one-tailed *t-*test, *p* = 0.0433), indicating that the staining capacity may be less efficient than anticipated (**Supplementary Fig. 6b**). Furthermore, the size of the EVs by TEM quantification only slightly albeit insignificantly correlated with the fluorescence intensity of the EVs by TIRFM quantification (Pearson’s correlation coefficient, *p* = 0.52 for *r* = 0.12), indicating that fluorescence intensities from TIRFM images are likely due to expression and not size (**Supplementary Fig. 6c**). TEM images further revealed that single fluorescent signals may originate from two neighboring EVs (**Supplementary Fig. 6d**), highlighting the limitations of diffraction-limited techniques, such as TIRFM, and underscoring the importance of the NPMx technique in correlating fluorescent signals to immunogold sorted EVs. Although the TRPS distributions for the bioreactor-generated EVs and the fluorescent EVs demonstrated similar sizes, the ∼11-nm sized EP subpopulation was lost and the distribution better resembled the TRPS data for both fluorescent and non-fluorescent EVs (**Fig. 2e**), providing insight into the bias of tetraspanin-based capturing.

## Discussion and Conclusions

The benefits of using the NPMx technique include reducing the required concentration of EPs for EM, quickening EM acquisition, and improving immunogold labeling, while providing various modalities, such as the non-biased electrostatic capture of EPs, population-based sorting of EPs, and orthogonal superimposition of optical and non-optical imaging at a single-EP resolution. Current EM techniques are susceptible to drying-induced artifacts^27^ and require high concentrations of EPs for standard EM and immunogold phenotyping.^6^ For patient-derived biofluids, high concentrations of EPs may be a luxury, and reducing concentrations and reproducibility is sought to provide usability across samples. The NPMx not only promotes the binding of EPs onto the mesh surface through electrostatics or user-defined immunocapture but also provides the precise coordinates of EPs on the surface, reducing acquisition time, decreasing required concentrations, and providing a possible future in high-throughput screening. While nanoparticles are innately subject to Brownian motion, leading to approaches to reduce the stochasticity of movement.^6,28-30^ We circumvented the natural phenomena via the immobilization of one particle species in a colloid mixture thus reducing the dimensionality of particle interactions. The engineered approach enhanced immunogold labeling and provided the precise coordinates of capture to easily superimpose TEM and TIRFM images and measurements for the first time.

While the proposed NPMx technique demonstrates a proof-of-concept to accelerate the characterization of EPs with state-of-the-art technology, we acknowledge some limitations to the technique. While advantageous, the NPMx technique requires lower concentrations of EPs to limit EP aggregation and reduce false-positive binding events. Furthermore, the resolution of SRM and EM images differ, leading to offsets and slight misalignment during image superimposition. Despite these limitations, the NPMx technique offers various modalities and combinatorial approaches that enable flexibility in experimental design, allowing for investigations for all types of EPs. The success of correlating optical and non-optical techniques is a step forward in reducing batch-to-batch variability by merging the best aspects of each technique for a comprehensive single-EP analysis.

## Author Contributions

E.R., K.T.N., and X.Y.R. developed the technology. E.R., K.T.N., and X.Y.R. prepared the figures and wrote the manuscript. C.L.H. prepared the bioreactor EVs and performed quality control. J.D.R. performed TRPS. C.K.N. developed the rotating algorithm. E.R. supervised the whole study. All authors provided critical feedback and helped to shape the research, analysis, and manuscript.

## Acknowledgment

Electron microscopy was performed at the Center for Electron Microscopy and Analysis (CEMAS) at The Ohio State University. This work was supported by the U.S. National Institutes of Health (NIH) and the National Center for Advancing Translational Sciences (NCATS) under grants UG3/UH3TR002884 (E.R.) and U18TR003807 (E.R.). The William G. Lowrie Department of Chemical and Biomolecular Engineering and the Comprehensive Cancer Center at The Ohio State University provided additional support for E.R.

