## Supplementary Information for "Limiting Brownian Motion to Enhance Immunogold Phenotyping and Superimpose Optical and Non-Optical Single-EP Analyses"

[†]Authors contributed equally

**Supplementary Table 1: List of antibodies.**

| Antibody | | | Catalog number | Supplier |
|---|---|---|---|---|
| Capture | EVs | Anti-CD63 | MAB5048 | R&D Systems |
| | | Anti-CD9 | MAB1880 | R&D Systems |
| | MLV-V5 | Anti-V5 tag | AB9116 | Abcam |
| Detection | EVs | Anti-CD63 (Alexa Fluor® 488) | SC-5275 AF488 | Santa Cruz Biotechnology |
| | MLV-V5 | Anti-V5-FITC | R963-25 | Thermo Fisher Scientific |

**Supplementary Table 2: Quantification of Strep-AuNP adsorption.**

| 0.25% w/v AuNP | Position | Count | Area (μm²) |
| --- | --- | --- | --- |
| 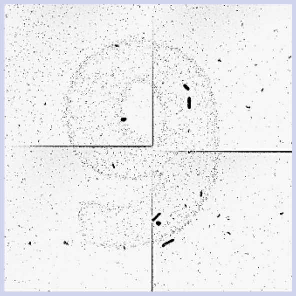 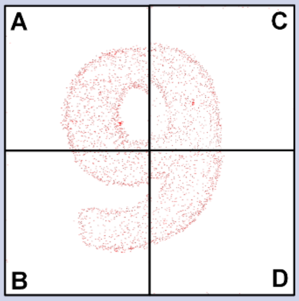  | 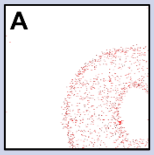  | 1229 | 1.82E+02 |
| | 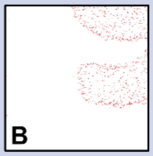  | 745 | 1.50E+02 |
| | 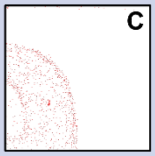  | 971 | 1.53E+02 |
| | 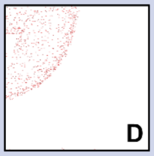  | 714 | 1.35E+02 |
| **0.5% w/v AuNP** | **Position** | **Count** | **Area (μm²)** |
| 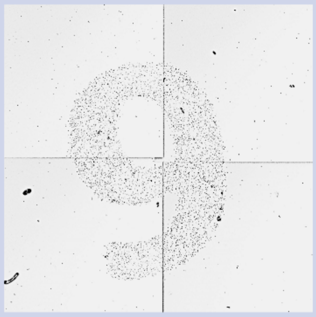 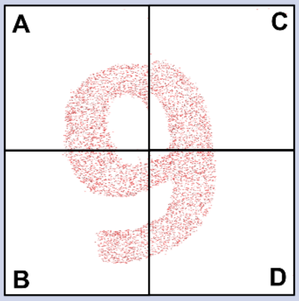  | 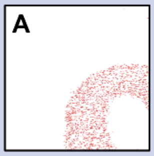  | 1664 | 1.63E+02 |
| | 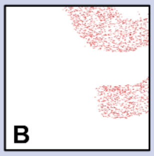  | 1655 | 1.61E+02 |
| | 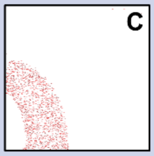  | 1420 | 1.27E+02 |
| | 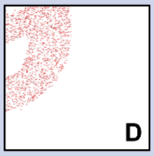  | 1769 | 1.68E+02 |

**Supplementary Table 3: Quantification of Strep-AuNP adsorption.**

| 0.25% w/v AuNP | Position | Count | Area (µm²) |
|---|---|---|---|
| 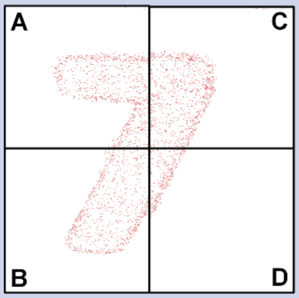 | 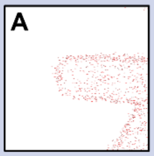 | 876 | 1.56E+02 |
| | 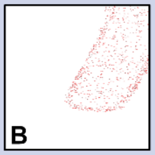 | 825 | 1.69E+02 |
| | 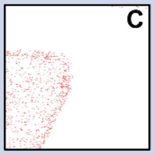 | 939 | 1.51E+02 |
| | 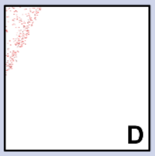 | 280 | 3.70E+01 |
| **0.5% w/v AuNP** | **Position** | **Count** | **Area (µm²)** |
| 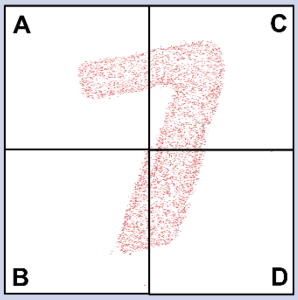 | 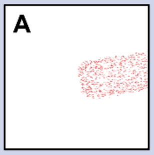 | 797 | 9.37E+01 |
| | 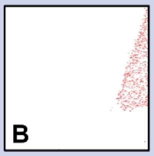 | 675 | 7.85E+01 |
| | 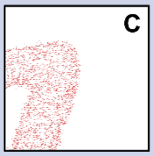 | 1931 | 2.14E+02 |
| | 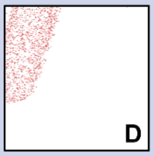 | 1280 | 1.27E+02 |

**Supplementary Table 4: Quantification of Strep-AuNP adsorption.**

| 0.25% w/v AuNP | Position | Count | Area (µm²) |
|---|---|---|---|
| 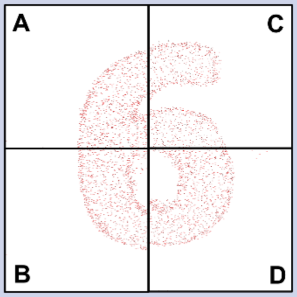 | 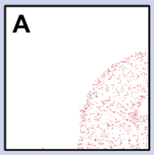 | 1000 | 1.20E+02 |
| | 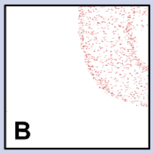 | 999 | 1.24E+02 |
| | 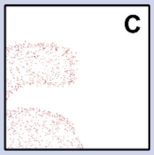 | 1012 | 1.33E+02 |
| | 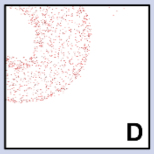 | 1104 | 1.43E+02 |
| **0.5% w/v AuNP** | **Position** | **Count** | **Area (µm²)** |
| 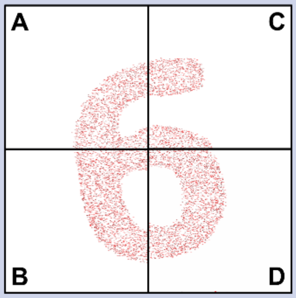 | 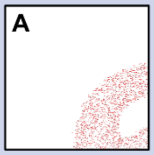 | 1593 | 1.20E+02 |
| | 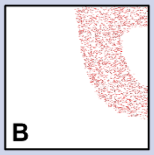 | 2160 | 1.54E+02 |
| | 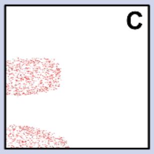 | 1223 | 8.64E+01 |
| | 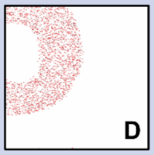 | 2312 | 1.59E+02 |

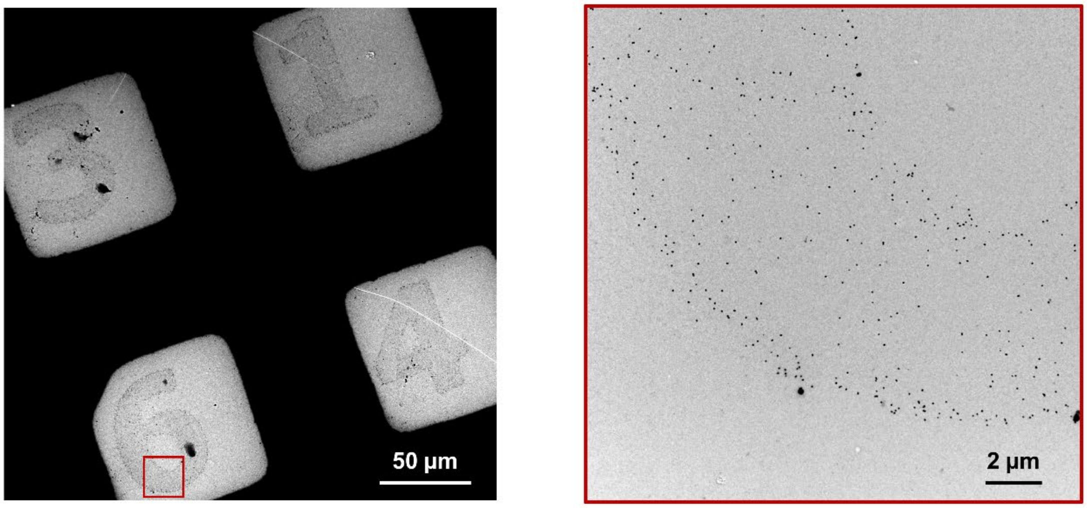


**Supplementary Figure 1: Single numeral Strep-AuNP micropatterns on individual square meshes.** Single numeral micropatterns consisting of Strep-AuNPs are adsorbed onto single square meshes on the transmission electron microscopy (TEM) grids. The square is magnified in the proceeding inset.

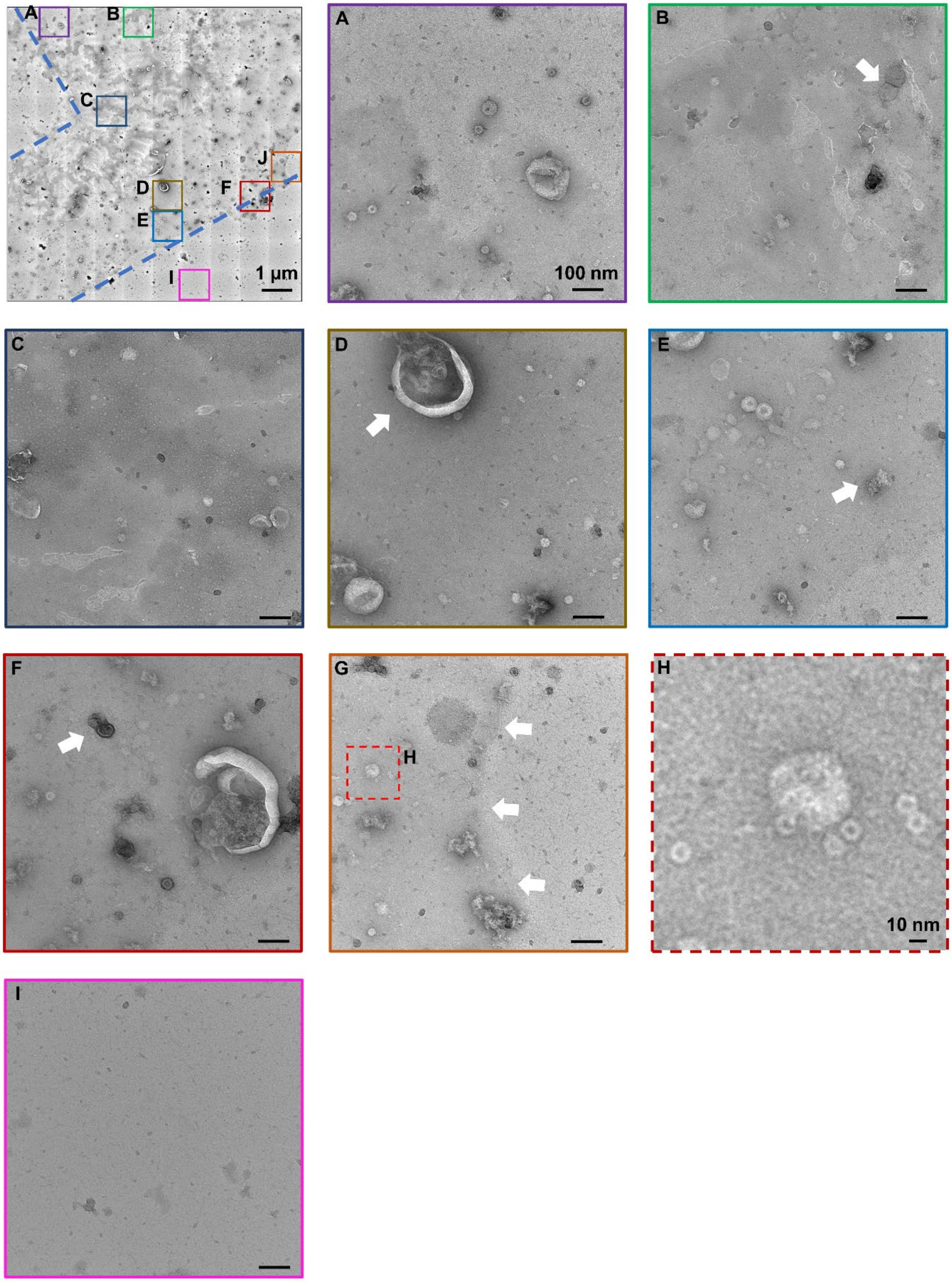
A
B
C
D
E
F
I
J
1 µm
A
100 nm
B
C
D
E
F
G
H
H
10 nm
I

**Supplementary Figure 2: Electrostatic immobilization of EVs onto a micropattern.** The electrostatic immobilization of extracellular vesicles (EVs) enables the non-biased capture of EVs onto the mesh surface. The squares are magnified in the proceeding inset, where **(A)** illustrates the capture of EVs onto the polycation surface with a classical "cup shape" morphology, **(B)** electron-dense EVs without the classical "cup shape" morphology, **(C)** the polycation surface mostly devoid of EVs, **(D)** multicompartment EVs, **(E)** protein aggregates, **(F)** positively stained EVs with compromised membranes, **(G)** the boundary of the polycation surface, **(H)** magnification of the square in **G** with minuscule EP structures, and **(I)** the mesh outside the functionalized micropattern region. The arrows indicate the location of the aforementioned phenomena.

a)

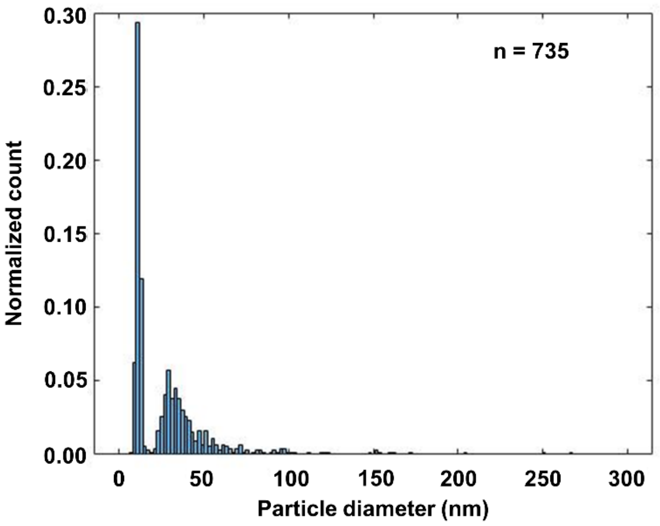
n = 735
Normalized count
0.30
0.25
0.20
0.15
0.10
0.05
0.00
0
50
100
150
200
250
300
Particle diameter (nm)

b)

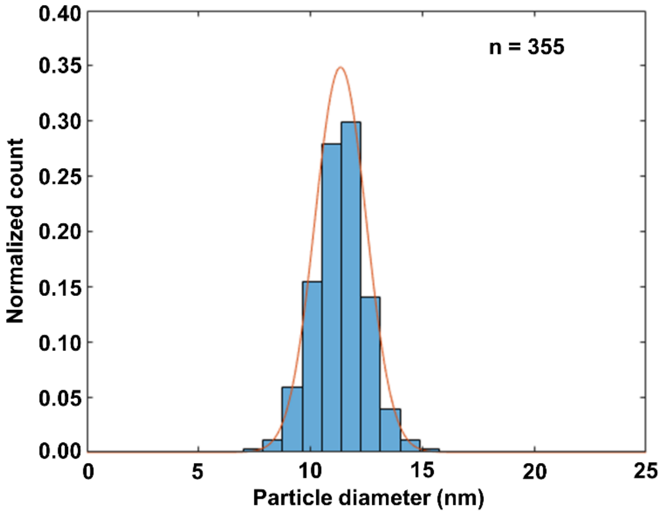
n = 355
Normalized count
0.40
0.35
0.30
0.25
0.20
0.15
0.10
0.05
0.00
0
5
10
15
20
25
Particle diameter (nm)

c)

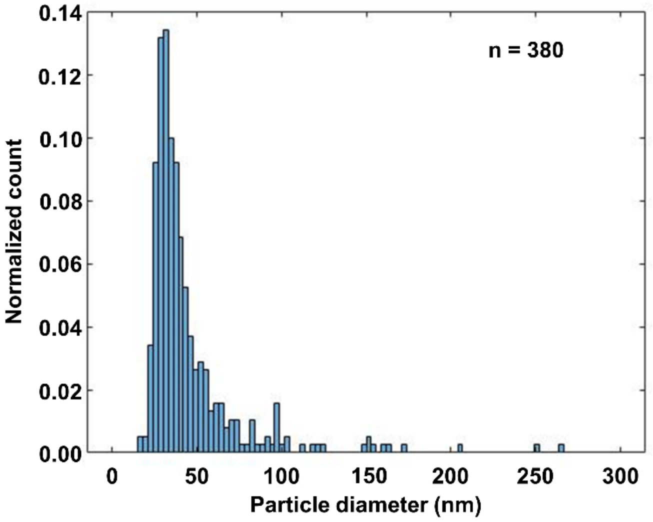
n = 380
Normalized count
0.14
0.12
0.10
0.08
0.06
0.04
0.02
0.00
0
50
100
150
200
250
300
Particle diameter (nm)

**Supplementary Figure 3: Quantification of EV size based on TEM images. (a)** The whole EV population demonstrates a bimodal distribution. **(b)** The smallest distribution is normally distributed and corresponds to the minuscule EP structures. **(c)** The largest distribution is left-skewed with the majority of particles falling below the detection limit of tunable resistive pulse sensing (TRPS).

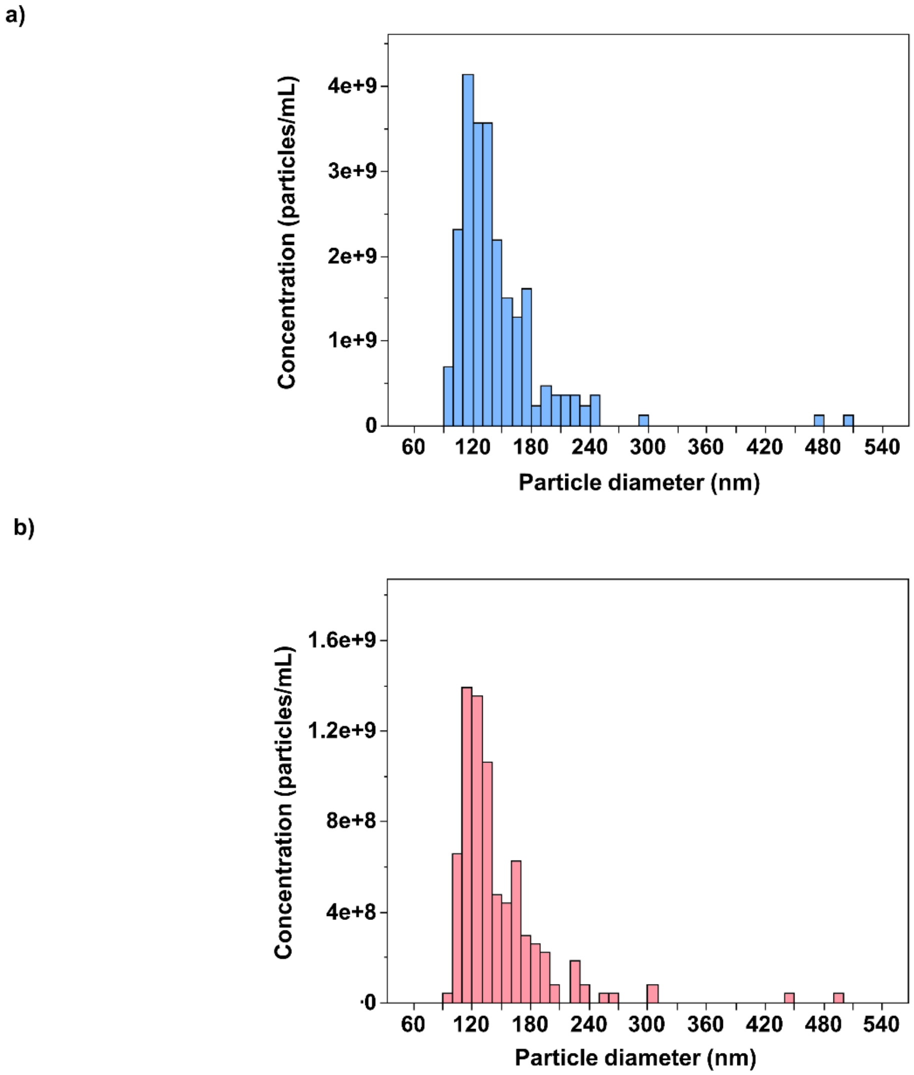


**Supplementary Figure 4: TRPS measurements. (a)** The TRPS measurements of the bioreactor-generated EVs do not contain the EV subpopulations measured in the TEM images. **(b)** The TRPS measurements of fluorescent EVs are similar in size to the bioreactor-generated EVs.

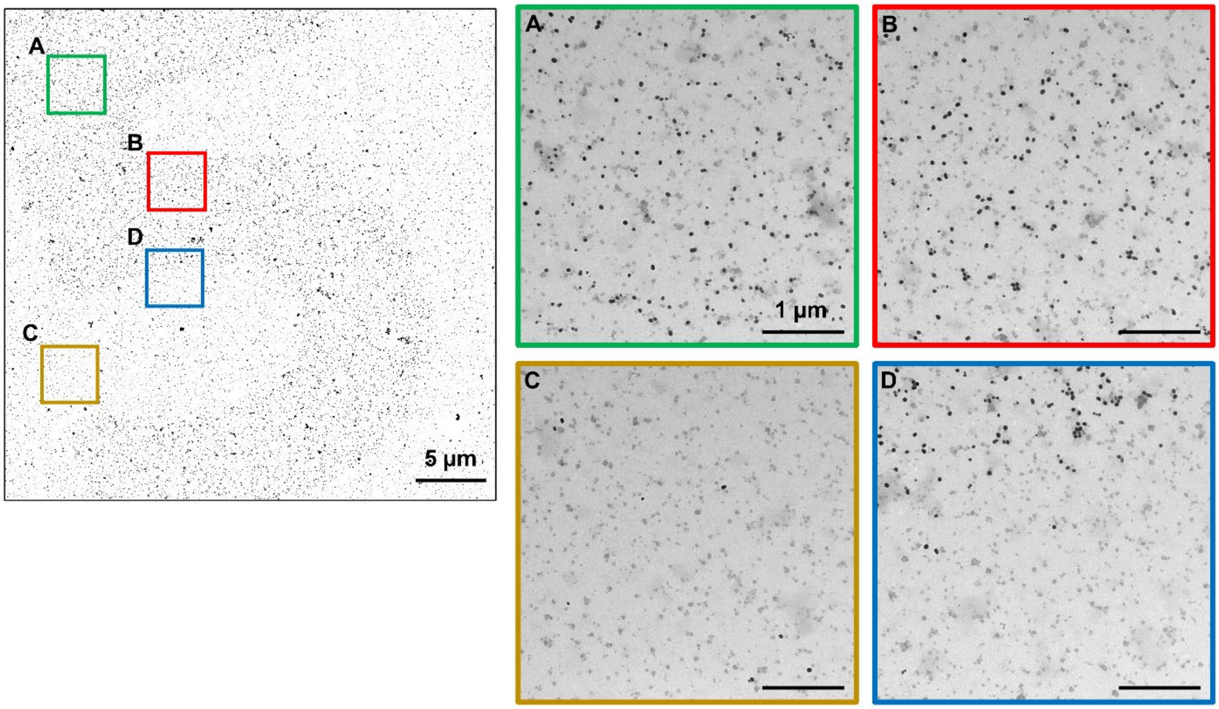


**Supplementary Figure 5: Negative control for the capture of EVs.** Strep-AuNPs without antibody functionalization are unable to capture EVs onto the micropatterned surface. The squares are magnified in the proceeding inset.

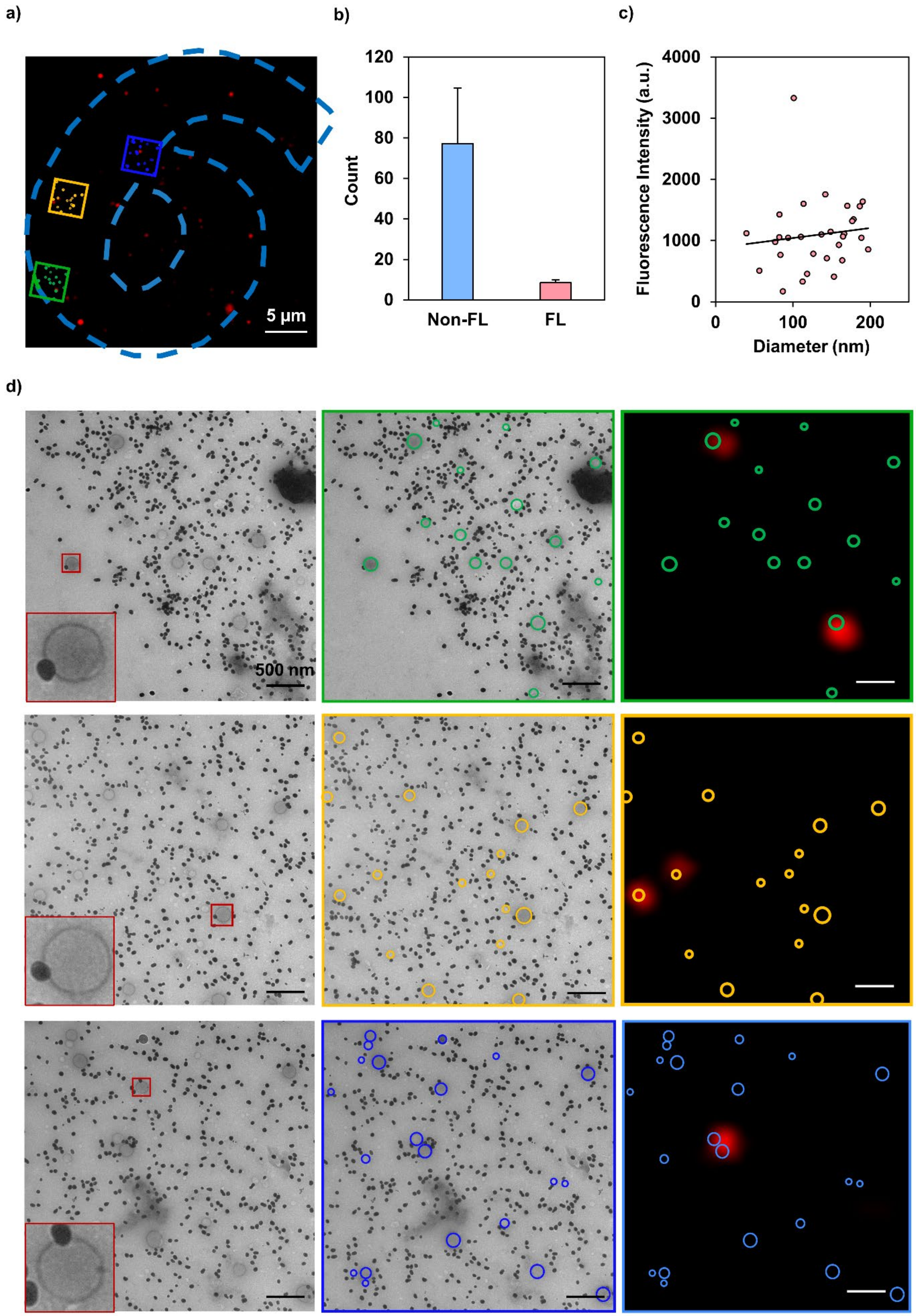

a)
b)
c)
d)
5 µm
Count
Non-FL
FL
Fluorescence Intensity (a.u.)
Diameter (nm)
500 nm

**Supplementary Figure 6: Superimposition of fluorescent EVs. (a)** Total internal reflection fluorescence microscopy (TIRFM) illustrates the specific capture of EVs within the micropatterned grid surface. The dotted outlines represent the boundary of the micropattern. **(b)** One-to-one mapping of EVs from TIRFM images to TEM images reveals that a majority of EVs are non-fluorescent (non-FL) and opposed to FL (N = 4 numerals, n = 3 fields of view; the error bars indicate the standard error of the mean). **(c)** The fluorescence intensity of EVs only correlates minimally with their measured diameter (N = 4 numerals, n = 3 fields of view). **(d)** The magnifications of the squares in **Supplementary Fig. 6a** are provided. The squares are magnified as an inset and the circles define the perimeters of EVs.
